# ATM-mediated DNA damage response in macrophages primes phagocytosis and immune checkpoint regulation

**DOI:** 10.1101/2020.03.14.987438

**Authors:** Aakanksha Bansal, René Neuhaus, Elena Izquierdo-Alvarez, Daniela Vorholt, Henning Feldkötter, Hendrik Nolte, Philipp Lohneis, Reinhard Büttner, Markus Krüger, Christian P. Pallasch, Björn Schumacher

## Abstract

Genotoxin-based cancer therapies trigger a DNA damage response (DDR) to eliminate cancer cells but similarly exert profound alterations to the immune cell compartment. Macrophages in the tumor microenvironment are important and multifaceted components in actively sustaining tumor progression but also in clearance of tumor cells and play an important role in the outcome of chemotherapy. Here, we report that post neo-adjuvant chemotherapy macrophages in tumor samples display elevated expression of the immune checkpoint PD-L1. We determined that chemotherapy or direct application of DNA damage to macrophages *in vitro* triggers a specific M_DDR_ polarization phenotype characterized by increased phagocytic activity, elevated PD-L1 expression, and induction of endotoxin tolerance. Phospho-proteomics analysis revealed epigenetic alterations and ATM-dependent induction of senescence in macrophages upon DNA damage. We propose that ATM-induced senescence and PD-L1 immune checkpoint induction mediate a specific M_DDR_ macrophage polarization that might contribute to chemotherapy responses. Furthermore, our results clarify the functional role of tumor associated macrophages in the context of DNA damage and combination therapies including checkpoint inhibition.

## Introduction

DNA damaging chemotherapy and radiation therapy is employed to eliminate malignant cells through apoptotic outcomes of the DNA damage response (DDR). However, non-transformed components of the tumor microenvironment also play an important role in the therapeutic outcome and adverse events (1). Systemic chemotherapy is affecting multiple tissues and cell types. In particular, hematotoxicity represents a major proportion of adverse events in genotoxic therapies with effects on immune cells showing a multitude of effects ranging from T-cell depletion and subsequent immune suppression to complex inflammatory responses.

Within the tumor microenvironment macrophages have emerged as key regulators of disease progression and immune regulation. Moreover, macrophages are highly plastic phagocytic cells, capable of performing tissue-specific homeostatic, protective and pathogenic functions (2,3). Macrophages show plasticity in their programming and are capable of switching from one functional phenotype to another in response to variable microenvironment signals (e.g., microbial products, damaged cells, activated lymphocytes). They acquire context-dependent phenotypes that either promote or inhibit host antimicrobial defense, antitumor immunity and tissue repair (4,5). Macrophages represent a spectrum of activated phenotypes but are categorized in discrete polarized population such as, classically activated macrophages (M1), alternatively activated macrophages (M2) and tumor-associated macrophages (TAMs). M1 mount an innate immune response against a variety of bacteria, protozoa and viruses, and have roles in antitumor immunity, while M2 have anti-inflammatory functions and regulate wound healing. In contrast to M1, TAMs suppress antitumor immunity (6,7).

Compelling evidence suggests the interplay and overlap between the DDR and innate immune responses: Some pathogens can induce genotoxic stress (8,9), while macrophages generate potent genotoxic species – ROS and NOS, during infections and tissue injury (10). Notably, the DDR has been reported to be involved in diverse macrophage functions such as inflammatory and antimicrobial responses and in the protection against sepsis (11–15). Specifically, ROS plays a role in differentiation of alternatively activated macrophages, while NOX2-induced ATM activation causes pro-inflammatory macrophage activation (13). Additionally, chemo- and radiotherapy induces DNA lesions leading to a DDR that consequently triggers the apoptotic demise or senescence of cancer cells (16). Recently, it has been suggested that in numerous chemotherapy regimens the induction of innate and adaptive immune responses towards malignant cells significantly contribute to anti-tumor outcomes (1,17,18). Radiation-induced changes in macrophages include M2-like polarization and have been associated with disease progression (19–21). Chemotherapy also increases the immunogenicity of malignant cells and disrupts immunosuppressive circuitries in the tumor microenvironment (1,22). DNA damage also leads to activation of immune responses and tumor cell death may impact immune cells in the tumor microenvironment (23). We have recently reported that cyclophosphamide treatment rewires the tumor microenvironment thus enabling macrophages to exert effector functions against lymphoma cells in the context of antibody-mediated chemoimmunotherapy via phagocytosis of tumor cells (24).

Despite a plethora of studies linking the DDR with macrophages, the reprogramming of macrophages in response to DNA damage remains incompletely understood (25). Here we report clinical evidence of specific alterations in macrophages induced by DNA damage. We report that the DDR primes macrophages, thus altering their functional orientation. We characterized the functional phenotype of DNA damage-primed macrophages and through a phospho-proteomics approach reveal a comprehensive map of post-translational modifications of ATM target proteins. DNA damage-priming of macrophages caused a fundamental phenotypic shift, involving changes in histone organization and modification, metabolic pathways, followed by altered surface receptor repertoire and functional reprograming. Collectively, our findings show that the ATM-mediated DDR primes macrophage and modulate its immune function.

## Results

### Chemotherapy- induced DNA damage modulates Macrophage function

We primarily analyzed the effects of neo-adjuvant chemotherapy on macrophages in follow-up samples in three cohorts of sarcoma, esophageal and breast cancer patients. Comparing biopsy specimens prior to neo-adjuvant chemotherapy with surgery tumor specimens post chemotherapy, we detected a significant increase of PD-L1 expression in CD68 positive macrophages by immunohistochemistry (Figure 1A, Supplemental Figure 1) (McNemar test p=0.044, Chi-square 8.1). Furthermore, we observed a change in morphology of macrophages after chemotherapy and an increase of phagocytosis in post-chemotherapy tissues (Figure 1B). These findings indicate a functional shift in tumor-associated macrophages after chemotherapy. Since we have observed rewiring of macrophage compartments in the context of alkylating DDR via an indirect cytokine response from lymphoma cells (26), we now intended to further dissect the mechanisms of DNA damage-induced functional changes. In order to determine the immediate effects of DNA damage in macrophages, we further addressed the DDR in macrophages *ex vivo* to exclude tissue dependent indirect modulation such as paracrine effects.

**Figure 1:**
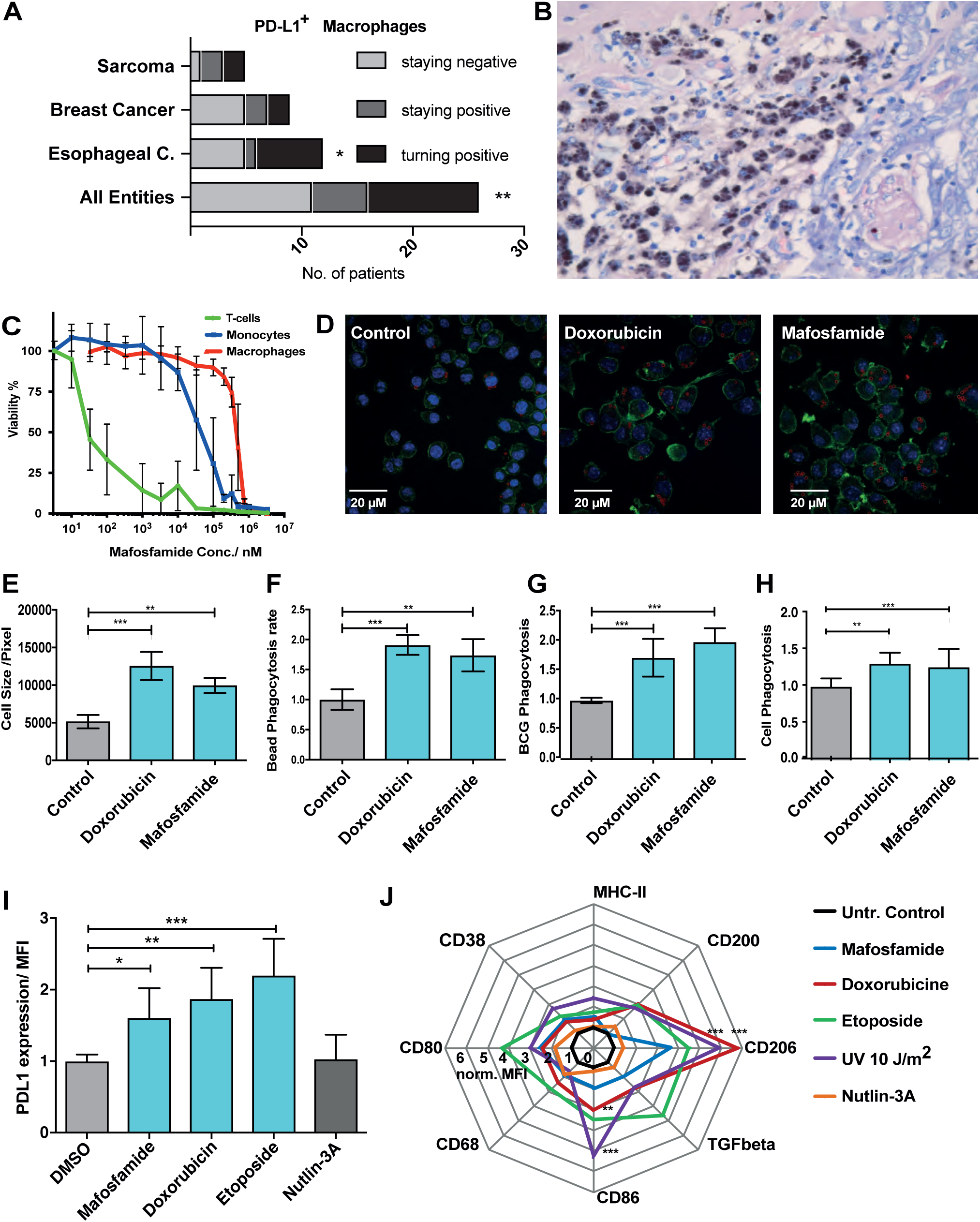
DNA damage modulates macrophage phenotype and increases phagocytic acitivity. A) A bar graph displaying PD-L1 status of CD68+ macrophages in cancer patient samples prior and post chemotherapy B) A micrograph example of breast cancer tissue (Giemsa) after neoadjuvant treatment displaying multiple macrophage with high phagocytic activity C) Dose response curve of in vitro treatment human primary T-cells, monocytes and macrophages D) Example micrographs of macrophage in vitro treatment with Doxorubicine and mafosfamide with assessment of phagocytosis activity of fluorescent beads (red) E) a bar graph displaying significantly increased macrophage cell size after chemotherapy treatment in vitro, F) a bar graph showing increased phagocytic bead uptake after chemotherapy treatment G) a bar graph showing increased phagocytic BCG-mycobacteria uptake after chemotherapy treatment H) a bar graph showing increased antibody-dependent cellular phagocytosis (ADCP) of hMB-lymphoma cells after chemotherapy pre-treatment of macrophages I) a bar graph displaying increased PD-L1 expression after in vitro treatment J) a radar plot displaying macrophage marker expression profile, mean fluorescence intensity rate normalized to untreated control (values represent the mean ± SD of three biological replicates; * = p ≤ 0.05, ** = p ≤ 0.01, *** = p ≤ 0,001).

To assess the direct effects of chemotherapy on macrophage viability, we addressed induction of apoptosis by chemotherapy in human monocyte-derived macrophages *in vitro*. DNA-alkylating treatment by mafosfamide (as an *in vitro* surrogate for cyclophosphamide therapy) induced apoptosis in very high dosages (IC50 46 µM) in macrophages, whereas monocytes (IC50 3.9 µM) and particularly T-cells (IC50 1.8 nM) displayed higher susceptibility towards induction of apoptosis by mafosfamide (Figure 1C). Chemotherapy exerts a plethora of cell-autonomous and non-autonomous effects. To differentiate the observed changes in macrophage phenotype after chemotherapy being induced as an indirect response to tissue damage or a direct effect mediated by the DDR, we next analyzed various macrophage cell lines and primary macrophages *in vitro*. In J774A.1 macrophages treated with mafosfamide or doxorubicin we primarily observed a change in cellular morphology with enlarged cell size, increased cytoplasmic volume and a significant increase in phagocytosis of fluorescently labeled beads (Figure 1D-F). Phagocytosis of fluorescently labeled BCG mycobacteria was similarly elevated significantly after mafosfamide and doxorubicine treatment (Figure 1G). Moreover, we assessed the antibody-dependent cellular phagocytosis (ADCP) of lymphoma cells by the anti-CD52 antibody alemtuzumab (Figure 1H). Here, pre-treatment of macrophages revealed significantly increased tumor cell clearance. Similar to the post-chemotherapy patient samples, we observed a direct induction of the immune checkpoint PD-L1 on macrophages by mafosfamide, doxorubicin and etoposide. The MDM2 inhibitor and indirect p53 activator Nutlin-3A, however, did not induce PD-L1 expression (Figure 1I). To further characterize the immunophenotype of macrophages following DNA damage we assessed markers of macrophage polarization and observed particularly increased expression of the M2-marker CD206 (Figure 1J) (27) as well as increased expression of M1-polarization markers CD68, CD80 and CD86. In contrast, the MDM2 inhibitor Nutlin-3A did only induce insignificant changes in polarization markers expression.

CD38, which was shown to be induced on macrophages in response to the senescence associated secretory phenotype (SASP) (28), was also induced after DNA-damage. As such, upon DNA damage, we observe a CD86^high^, CD206^high^, CD38^high^ PD-L1^high^ immunophenotype of the macrophages that does not display a pure M1- or M2-polarization

In conclusion, upon DNA damage-inducing chemotherapeutic drug exposure macrophages exhibit a phenotype that is characterized by altered expression of immune checkpoints, morphological changes, and increased phagocytic function.

### Macrophage priming with UV-induced DNA damage modulates its functional plasticity

While chemotherapy might display a combination of pure DNA damage and other compound-mediated effects, UV irradiation provides a most pure application of DNA damage. We employed UVC irradiation because treatment with 254 nm results in clearly defined physically-induced cyclobutane pyrimidine dimer (CPD) and pyrimidine 6-4 pyrimidone (6-4PP) DNA lesions. CPDs and 6-4PPs serve as experimental paradigm for helix-distorting lesions, which block replicative DNA polymerases as well as elongation of RNA polymerases. In addition, UV-induced lesions can be precisely dosed and chromatin re-organization after UVC-induced DNA damage is well-studied (29–34). Therefore, we employed UV-induced DNA damage as experimental cause-effect paradigm for genotoxins that similarly to chemotherapeutic agents interfere with replication and transcription elongation. Henceforth, UV-primed macrophages are referred as “M_DDR_” and mock-treated macrophages as “M_C_”.

Similar to chemotherapeutics-primed macrophages, M_DDR_ macrophages showed a modulation of macrophage polarization markers (Figure 1J), significant upregulation of CD206 (Figure 2A) and PD-L1 (Figure 2B) compared to M_C_ macrophages. We also observed morphological changes such as increased cell size and enhanced F-actin staining in M_DDR_ macrophages (Figure 2C).

**Figure 2:**
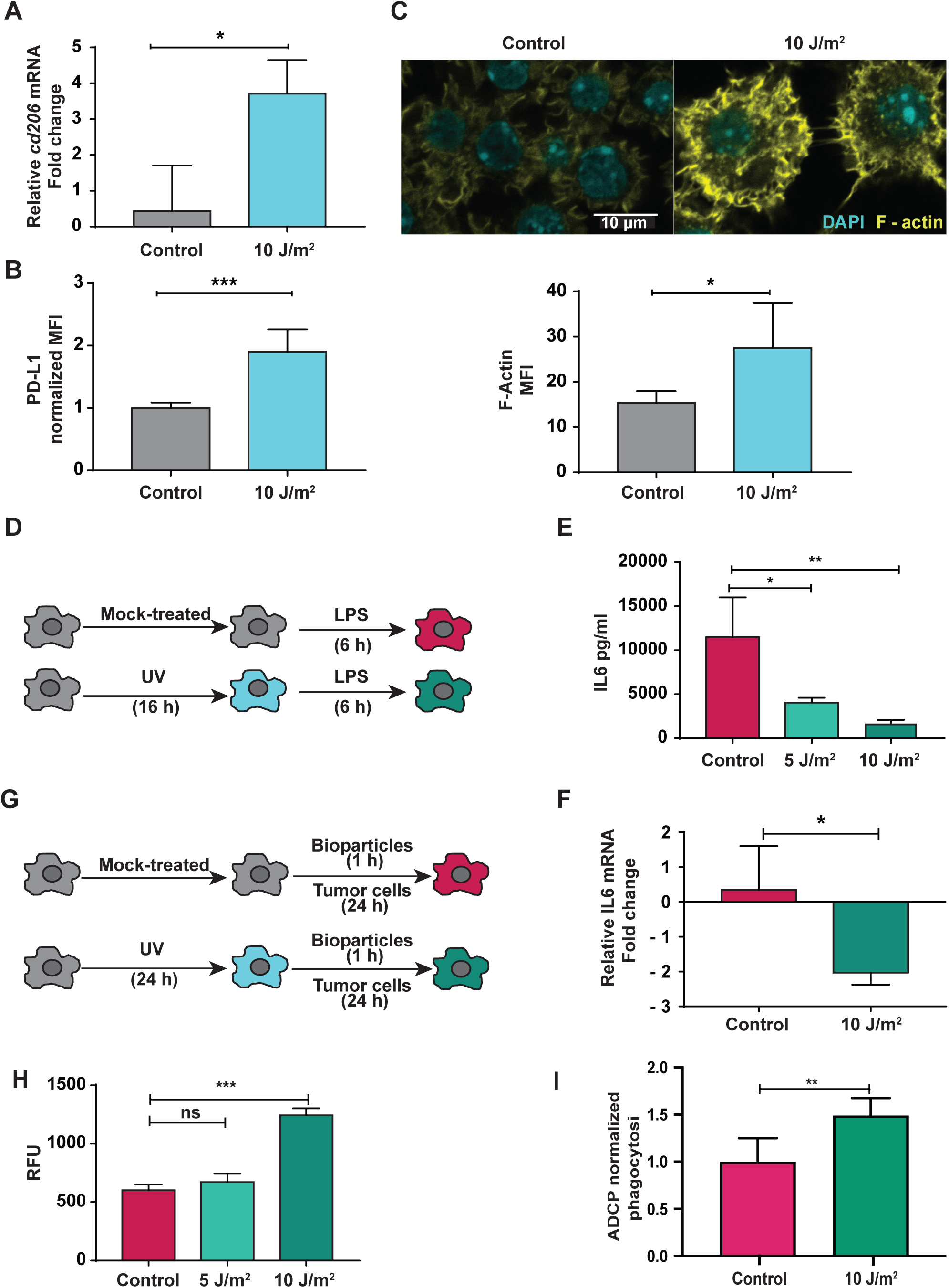
DNA damage modulates macrophage inflammatory response to LPS and phagocytosis: J774A.1 Macrophages were treated with UVC or chemotherapeutics and allowed to recover for 24 h and RT-qPCR and FACS was performed. A) M_DDR_ have higher fold change of cd206, shown relative to the expression in M_C_. RT-PCR quantification of mRNA extracted from M_DDR_ and M_C_ 24 h after treatment, normalized to beta-Tubulin and Peptidyl-prolyl cis-trans isomerasa. B) FACS analysis with antibody against PD-L1 or isotype controls shows more PD-L1 expression in M_DDR_ compared to M_C_. C) Immunofluorescence staining for F-actin(yellow), and DAPI (blue) of the M_DDR_ and M_C_ post 24 h of UV exposure. D,G) Working scheme for functional assays – endotoxin tolerance and phagocytic capacity of macrophages post direct DNA damage induced by exposure to UV. E) ELISA for IL6 quantification, supernatant collected from M_DDR_ and M_C_ macrophages at 0 h after LPS exposure. F) RT-PCR of *IL6* mRNA extracted at 0 h after LPS exposure quantified over M_C_ macrophages, normalized to ***60S acidic ribosomal protein P0*** (*<u>Rplp0</u>*) and *Peptidyl-prolyl cis-trans isomerase* (*<u>Ppia</u>*). H) Relative fluorescence unit (RFU) is measured 1 h after M_DDR_ and M_C_ incubation with pHrodo E.coli. (values represent the mean ± SD of three biological replicates \**P* < 0.05, \*\**P* ≤ 0.01 and \*\*\**P* ≤ 0.001 (unpaired two-sided Student’+s *t*-test).

We next assessed the functional plasticity of macrophages upon UV-induced DNA damage. We employed two *in vitro* assays to study different aspect of M_DDR_ macrophage function – (i.) response to inflammatory stimuli (Endotoxin tolerance (ET) assay) (Figure 2D) and (ii.) phagocytosis (Figure 2G). In a classical ET assay, macrophages that have been pre-challenged with LPS (primed) secrete less inflammatory cytokines - IL6 (mouse) and TNFα (human THP-1 cells) (35) when they are re-challenged with LPS compared to unprimed macrophages. To determine whether DNA damage modulates macrophage function, LPS pre-treatment was replaced with a low non-lethal UVC dose, and macrophages were examined for induction of the endotoxin tolerance phenotype.

Strikingly, the level of IL6 secretion and *IL6* mRNA both were strongly reduced in response to the LPS challenge in UV-primed J774A.1 macrophages compared to naïve J774A.1 macrophages (Figure 2E, F). Similarly, the level of IL6 secretion was also reduced in response to LPS challenge in UV-primed RAW264.7 (Figure S2A) and peritoneal macrophages (Figure S2C). In UV-primed THP-1 macrophages, TNFα secretion was reduced (Figure S2B). The chemotherapeutics-primed macrophages also showed a trend towards downregulation of IL6 secretion upon subsequent exposure to LPS (Figure S2D). These findings suggest that UV-induced DDR mimics the LPS priming effect, resulting in a gain of endotoxin tolerance phenotype in M_DDR_ macrophages, similar to LPS re-challenged macrophages.

We next analyzed antibody independent phagocytosis capacity by incubating M_DDR_ macrophages and M_C_ macrophages with pHrodo *E.coli* bioparticles. These bioparticles are non-fluorescent outside of cells but fluoresce bright red in phagosomes. Strikingly, M_DDR_ macrophages exhibit higher relative fluorescence units (RFU) compared to M_C_ macrophages (Figure 2H, Figure S2E). UV-irradiated macrophages also showed significantly enhanced capability to provide ADCP of lymphoma target cells (Figure 2I).

Collectively, our findings show that DNA damage primes M_DDR_ macrophages to attain an endotoxin tolerance phenotype and modulate their immune function towards increased phagocytosis and regulation of the adaptive immune response via up-regulation of the immune checkpoint PD-L1.

### Phosphoproteomics analysis indicates chromatin remodeling and a central role of ATM signaling in the macrophage DDR

To gain further insight into the molecular mechanisms underlying the DDR in macrophages, we performed phosphoproteomics of UV-primed macrophages. Phosphopeptides were generated by in-solution digestion followed by TiO_2_ bead-based enrichment. Phosphorylated sites were localized and quantified using a label-free based liquid chromatography and tandem mass spectrometry (LC-MS/MS) approach (Figure 3A).

**Figure 3.**
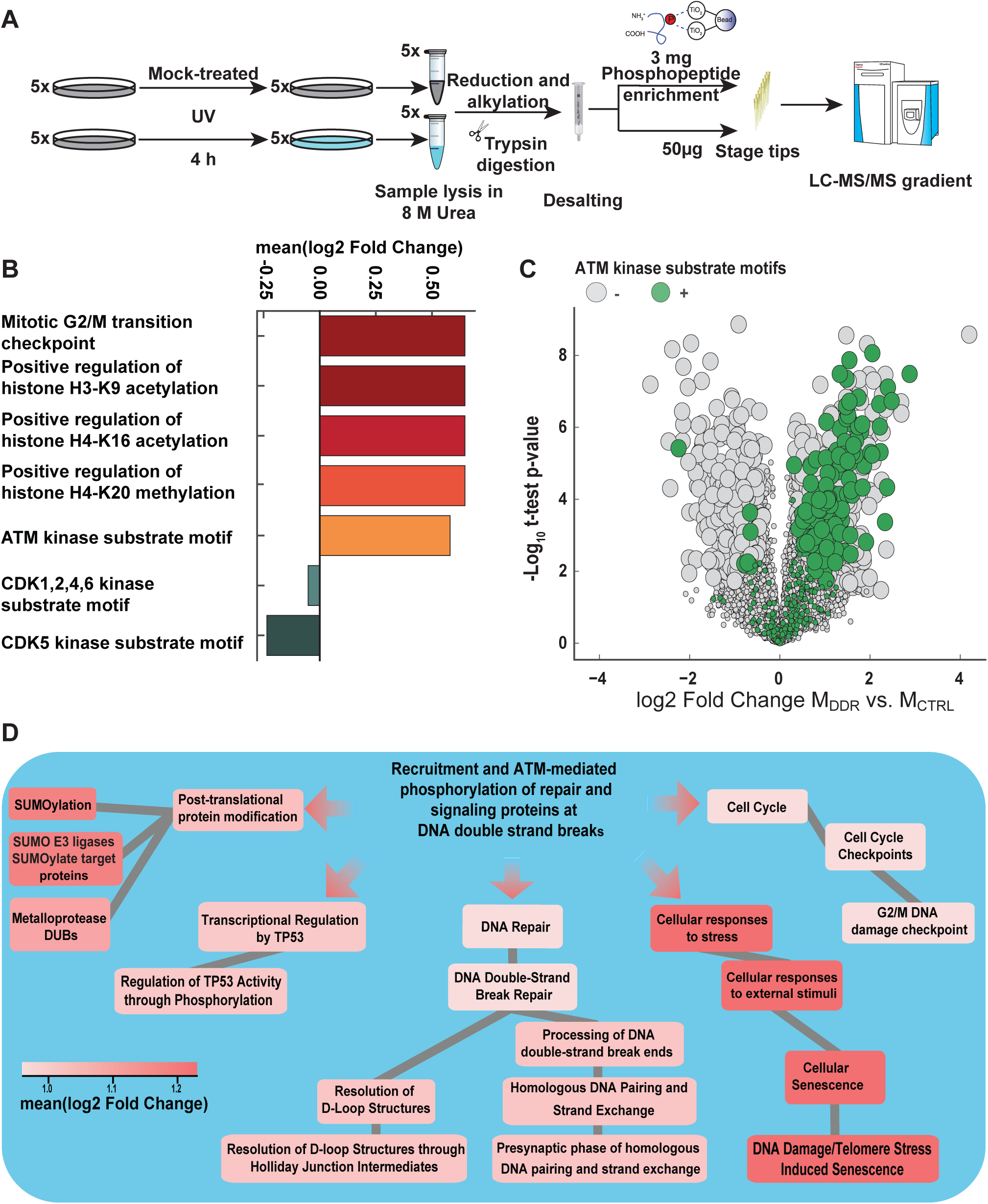
Phosphoproteomics analysis of M_DDR_. A) Experimental workflow: J774A.1 were mock-treated or UVC (10 J/m2) treated. Post 4 h of treatment sample were collected and the phosphoproteome was analyzed by LC-MS/MS. B) Significant GO and motif terms representation identified via an 1D enrichment approach. C) Volcano plot of phosphosites detected upon UVC treatment. Significantly (FDR < 0.01) different phosphosites are highlighted with big circles and ATM kinase motif containing phosphosites are highlighted by green color. D) Reactome network for ATM-mediated phosphorylation using the phosphoproteomics data from M_DDR_ and M_C_. The color is based on the mean Log2FC value of UV compared to Control (FDR <0.01). Dark red represents the maximum mean Log2FC =1.5, and light red represents the minimum mean Log2FC =0.9.

To exclude that detected alterations of the phosphorylation peptides originated from different abundance of the corresponding protein, we measured the proteome using a data-independent approach. None of the 4969 detected proteins were significantly (FDR < 5%) changed in abundance (Figure S3A; Table S1) suggesting that the macrophage respond within the tested time frame to DNA damage primarily through post-translational modifications (PTMs). Therefore, phosphorylation peptide abundance was not normalized to the protein level.

We identified 9.160 phosphosites, out of which 1.305 phosphosites were significantly (FDR < 0.01) upregulated (65%) or downregulated (35%) in M_DDR_ compared to M_C_ (Figure S3E; Table S2). The false discovery rate was estimated by a permutation-based calculation using a fudge factor (s0) of 0.1 using a total number of 500 permutations (36). To obtain an unbiased systematic view of the phosphoproteome data, we performed a PCA analysis and found that the UV and control group segregated from each other (Figure S2B). Additionally, we performed correlation analysis and calculated the Pearson correlation coefficient between all experiments. (Figure S3C, D). Biological replicates of M_C_ and M_DDR_ clustered together indicating that the UV treatment induced unique phosphorylation patterns in macrophages.

To analyze the phosphoproteome changes systematically, we performed a 1D enrichment analysis of annotated Gene Ontology (GO), Kinase-Motifs, and Reactome terms (Table S3, S4). This type of analysis tests for every annotation term whether the corresponding log2 fold changes of the phosphorylation sites have a preference to be systematically up- or downregulated. A two-sample test is performed between the log2 Fold Change Distribution of the phosphorylation sites that were found to carry the ATM target consensus motif and all other remaining phosphorylation sites. The analysis provides a mean of the log2 fold changes between M_DDR_ and M_c_ of tested phosphorylation sites (here ATM target consensus motif) as well as an adjusted p value (FDR). The mean Log2FC value (t-test difference) obtained from 1D enrichment represents the mean of the Log2FC UV/control of all phosphosites of protein annotated by the respective GO terms, thus identifying systemic up or down regulation of categorical terms. The mitotic G2/M transition checkpoint associated phosphosites were highly induced suggesting a DNA damage checkpoint-induced cell cycle arrest in M_DDR_, while CDK1, 2, 4, 6, and CDK5 kinase substrate motifs were downregulated (Figure 3B).

Strikingly, phosphosites associated with the induction of histone H3-K9, H4-K16 acetylation as well as histone H4-K20 methylation were significantly increased. Among those, Brca1 phosphosites (P48754) were the only phosphosites that were commonly annotated in all of those three GO terms. Brca1 was significantly upregulated at 11 phosphosites (<u>S240</u>, <u>T522</u>, S710, S717, <u>S810</u>, <u>S815</u>, S831, S1481, <u>S1533</u>, S1584, and <u>S1585</u>). These phosphosites were matched to the PhosphoSite Plus^®^ database to identify the novel phosphosites, revealing 6 out of 11 that were not documented in the PhosphoSite Plus^®^ database (underlined). These data suggest that the DDR affect epigenetic modifications on histones.

The DNA damage-induced phosphorylation sites harboring the ATM target consensus motif Ser/Thr-Glu (S*/T*-Q) were the single most significantly enriched motif (p=9.04E-38) that was systematically upregulated (Mean Log2FC value (t-test difference)=0.58) in M_DDR_ compared to M_C_ (Figure 3C, 4). 129 of the significantly upregulated phosphosites sites were at the ATM specific consensus motif suggesting a central role of this DDR kinase (Figure 3C). We next generated a pathway network determining the overlap between functional annotations of ATM-target phosphorylation sites. As the “recruitment and ATM-mediated phosphorylation of repair and signaling proteins at DNA double-strand breaks” was significantly enriched, we analyzed its overlap with other GO terms with highly enriched phosphosites (Figure 3D).

We found the maximum overlap with the DNA damage/Telomere induced senescence and cellular senescence reactome pathway, which is consistent with the association of the physiological reprogramming of macrophages towards an M2-like phenotype with commonly accepted biomarkers of cellular senescence: constitutive *p16Ink4a* expression and senescence-associated β-galactosidase (SAβGal) (37).

Altogether, the GO term enrichment analysis suggests that post UV treatment, checkpoint activation occurs in M_DDR_ along with histone acetylation and methylation marks. Moreover, a very profound ATM kinase signature indicates that ATM might regulate the function of macrophages and promote a senescence phenotype in response to DNA damage.

### Phosphoproteome analysis reveals a regulatory network of M_DDR_

To generate an overview of phosphorylation sites that function within the detected reactome pathways, we created the network for over-represented pathways in the reactome database showing specific phosphorylation sites (Figure 4). ATM kinase (Q62388) hyperphosphorylated at S1987 was identified, which is used as a common biological marker for ATM kinase activation (38). In the phosphoproteomics data analysis, this was exemplified by hyperphosphorylation of ATM kinase targets such as γH2A.X (P27661) and NCL (P09405; S403, S425, <u>S563</u>).

**Figure 4.**
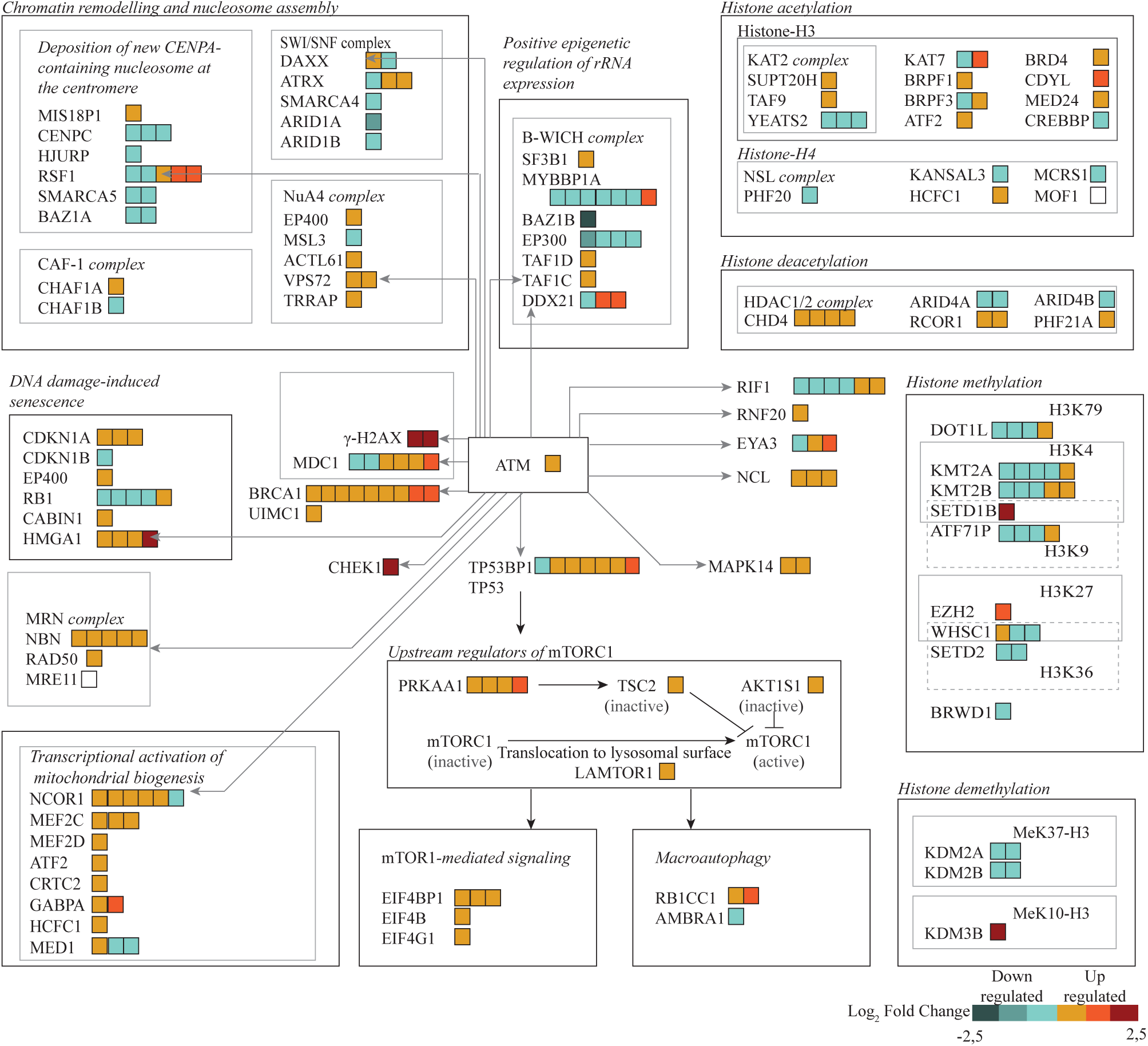
Network created via reactome analysis of phosphoproteome. The network for over-represented pathways identified via the reactome database showing specific phosphorylation sites upon UVC treatment. Symbols are as follows: filled rectangles, phosphopeptides detected by MS as downregulated (blue shades) or upregulated (red shades). Grey arrows indicated the phosphopeptides identified to have ATM kinase substrate motif in our phosphoproteomics.

The ATM-mediated γH2A.X forms the foundation of chromatin-based signaling cascade (39) involving histone PTMs, histone deposition and eviction, and nucleosome reorganization. γH2A.X binds directly to MDC1 (E9QK89; S176, T445, S592, S733, <u>S1437</u>, S1444) to further amplify the DDR. The histone chaperon NCL recruited via RAD50 (A8Y5I3; S574) (a subunit of MRN complex) (40) promotes nucleosome destabilization at DSBs. Interestingly, a study demonstrated that NBN (Q9R207; S398, S605, <u>S638</u>, <u>S644</u>, <u>S645</u>), which was highly phosphorylated in our analysis, a subunit of MRN complex plays a role in macrophage functional activity. Moreover, several studies reported that ATM is involved in diverse macrophage function, inflammatory response, antimicrobial response, and protection against sepsis (11–15).

We detected an expansive differential regulation of phosphoposites involved in histone modifications (Figure 4). Further coordination of DDR with histone dynamics occurs in the context of an ‘+Prime-Repair-Restore’+ model (31–34). According to this model, DNA damage is recognized, and histone-modifying proteins and chromatin remodelers reorganize local chromatin architecture to allow DNA repair factors to access and repair the DNA (41).

We identified two regulators of the onset of cellular senescence, hyperphosphorylated p53 (Q549C9, S389) and hypophosphorylated Rb (P13405) (active)(42). Strikingly, HMGA1 (P17095, S2, S6, S9, S44) is also hyper-phosphorylated in M_DDR_, which was reported to be an essential component of senescence-associated heterochromatin foci (SAHFs) (43). We furthermore identified phosphorylation of MAPK14 (B2KF34, S2, Y182) (also known as p38 MAP kinase) as a downstream target of ATM.

Taken together, the phosphoproteomics analysis suggests that ATM functions as the central regulator of the DDR in macrophages that impacts multitude of targets such as histone modifications, chromatin remodeling as well as p38 signaling.

### Epigenetic regulation of M_DDR_ phenotype induction

The pattern of epigenetic regulation in the phosphoproteomics appeared in a multitude of targets. In order to elucidate the functional relevance of this broad pattern we applied histone deacetylate inhibitor Trichostatin A (TSA) to inhibit heterochromatin foci formation. As the phosphoproteomics data indicate that the DDR in macrophages might induce senescence, we performed SAHF staining with DAPI and senescence-associated beta-galactosidasae (SA-β-Gal) staining in M_DDR_. SAHF foci formation and SA-β-Gal staining were significantly increased in M_DDR_ (Figure 5A, B). TSA treatment slightly reduced the number of SAHF positive cells but had no effect on SA-β-Gal staining and Annexin V staining (Figure 5A-D). We further assessed the endotoxin tolerance and phagocytosis capacity of M_DDR_ in the presence of TSA. TSA treatment reversed the reduced IL6 expression in response to LPS (Figure 5E) and significantly suppressed the phagocytosis capacity (Figure 5F) of M_DDR_ macrophages. Collectively, these data emphasize the role of DNA damage induced histone acetylation in modulation of macrophage function.

**Figure 5:**
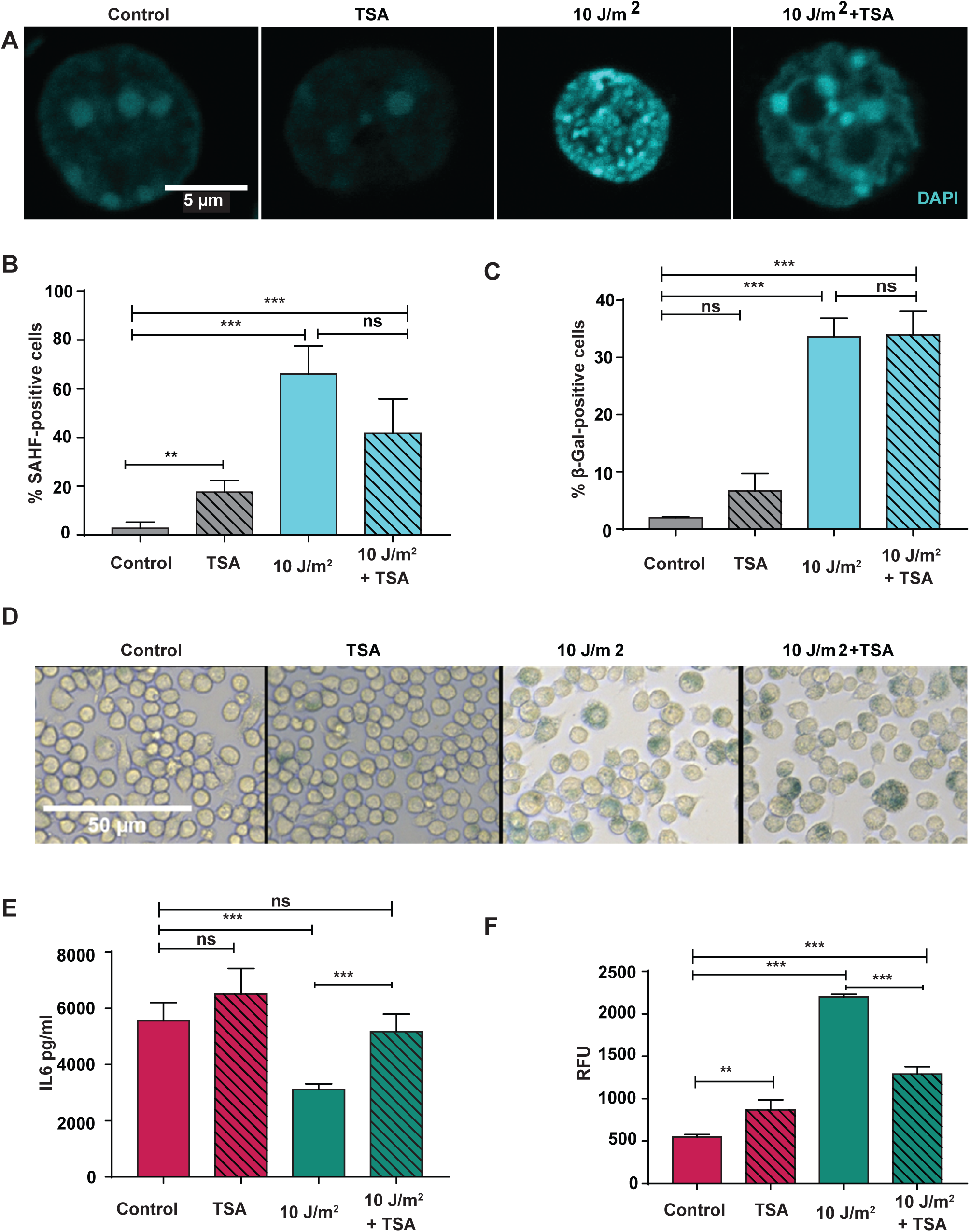
M_DDR_ show senescence phenotype and trichostatin A partially rescue M_DDR_ functional phenotypes. A, B) Immunofluorescence staining with DAPI 24 h of M_DDR_ with and without TSA and quantification. C,D) Βeta-Galactosidase staining of M_DDR_ with and without TSA. E) RT-PCR of IL6 mRNA extracted at 0 h of LPS treatment from M_DDR_ with and without TSA, quantified over M_C_ macrophages exposed to LPS, normalized to beta-Tubulin and Peptidyl-prolyl cis-trans isomerase. F) Phagocytosis assay of M_DDR_ with and without TSA. Values represent the mean ± SD of three biological replicates \**P* < 0.05, \*\**P* ≤ 0.01 and \*\*\**P* ≤ 0.001 (unpaired two-sided Student’+s *t*-test).

### ATM-dependent induction of M_DDR_ functional phenotypes

As the phosphoproteomics analysis showed ATM kinase substrate motifs containing phosphopeptides were highly upregulated upon exposure to DNA damage (Figure 3C,D; 4), we further examined whether ATM inhibition had any impact on SA-β-Gal staining. Remarkably, ATM inhibitor treatment not only diminished SA-β-Gal staining (Figure 6A,B) but instead induced apoptosis in M_DDR_ (Figure 6C). In line with this, Annexin V^+^ cells were detected in chemotherapeutic-treated macrophages in the presence of the ATM inhibitor (Figure S4A). We further assessed the endotoxin tolerance and phagocytosis capacity of M_DDR_ in the presence of the ATM inhibitor. ATM inhibition reversed the IL6 expression in response to LPS (Figure 6D) and also the phagocytosis capacity of M_DDR_ macrophages (Figure 6E). Moreover, the enhanced PD-L1 expression in M_DDR_ was also lost in presence of ATMi (Figure 6F).

**Figure 6:**
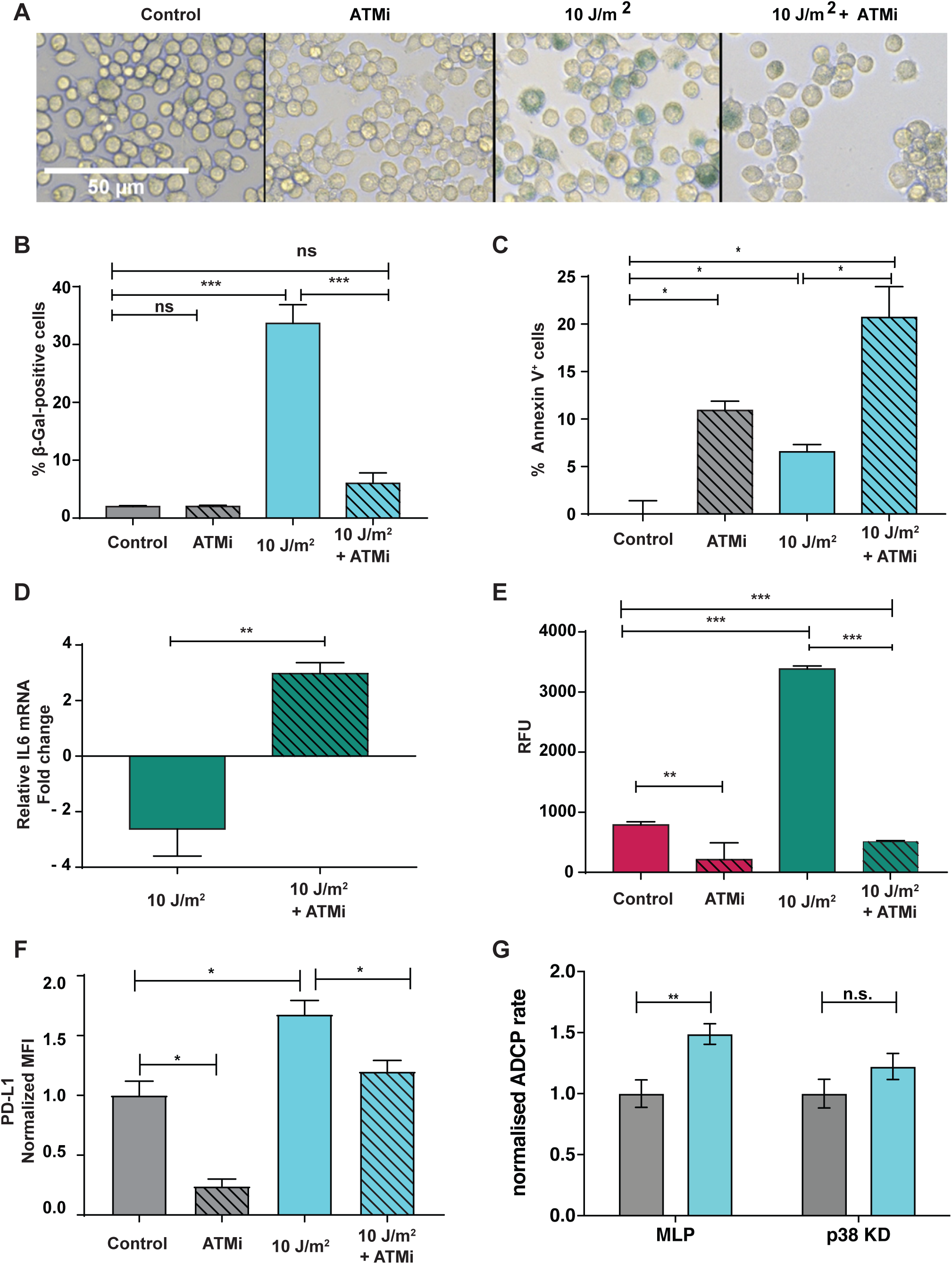
ATM inhibition reverses M_DDR_ functional polarization. A,B) Βeta-Galactosidase staining of M_DDR_ with and without KU 55933(ATMi). C) FACS analysis of M_DDR_ and M_C_ macrophages with and without KU 55933(ATMi) for Annexin V+ cells. D) RT-PCR of IL6 mRNA extracted 6 h after LPS treatment from M_DDR_ with and without KU 55933(ATMi), quantified over M_C_ macrophages exposed to LPS, normalized to beta-Tubulin and Peptidyl-prolyl cis-trans isomerase. E)) Phagocytosis assay of M_DDR_ with and without KU 55933(ATMi). F) FACS analysis with antibody against PD-L1 or isotype controls shows more PD-L1 expression in M_DDR_ compared to M_C_ with and without KU 55933(ATMi) G) ADCP of WT J774.1 macrophages vs. p38-targeted macrophages shows loss of UV-induced phagocytosis increase in p38-deficient macrophages Values represent the mean ± SD of three biological replicates \**P* < 0.05, \*\**P* ≤ 0.01 and \*\*\**P* ≤ 0.001 (unpaired two-sided Student’+s *t*-test).

To further elucidate phosphosites from our phosphoproteomic analysis downstream of ATM and dissect the relevance of particular pathways downstream of ATM. Here, we primarily elucidated the impact of p38 and p53 as major downstream components of the DNA damage checkpoint signaling in the regulation of key processes such as apoptosis and autophagy. We generated p38 and p53 deficient J774.A1 macrophages by specific shRNA retroviral transduction and used p38- and p53 deficient macrophages for functional analysis of DDR to UV and chemotherapy with mafosfamide and doxorubicin. We did not detect differences in PD-L1 expression due to p38- or p53-loss upon DDR treatment (Figure S4B). Assessing the capacity for ADCP, we also did not detect a decrease of phagocytosis in mafosfamide treated p53-deficient macrophages (Figure S4C). In contrast, p38 knock-down significantly diminished the phagocytic increase in comparison to empty vector control macrophages (Figure S4C). Similarly, p38 knockdown blunted the elevated phagocytosis in UV-treated macrophages (Figure 6G).

Collectively, these data show that genotoxic stress leads to phenotypic adaptations of macrophages through ATM mediating up-regulation of PD-L1, enhanced phagocytosis, endotoxin tolerance, cellular senescence and protection from apoptosis. Amongst the multiple ATM downstream effects histone modification particularly is involved in endotoxin tolerance and phagocytosis regulation, while p38 particularly regulates the phagocytic activity.

## Discussion

Our findings were initiated by observations in clinical samples showing that following clastogenic chemotherapy, macrophages display enhanced PD-L1 expression and increased phagocytosis. To define the mechanistic basis for the macrophage response to chemotherapy, we chose to examine the DDR in independent sources of macrophages utilizing both macrophage cell lines as well as primary peritoneal macrophages. To exclude any indirect effects of the chemotherapy such as cytokine release from the microenvironment and contact with apoptotic debris, we focused on the phenotypic alterations in macrophages upon direct infliction of defined DNA lesions through UVC treatment *in vitro*. We have shown here that the DDR in macrophages induces a specific M_DDR_ phenotype with increased cell size and elevated phagocytic activity. Particularly the expression of CD206 and PD-L1 indicate similarities of M_DDR_ consistent with M2 polarization.

We show that similarly to pre-challenge with LPS, macrophages build a non-specific memory upon being primed with DNA damage resulting in heightened phagocytosis and suppressed IL6 secretion when re-challenged with bioparticles and LPS, respectively. While the chronic DDR in many cell types such as fibroblasts triggers inflammation (44,45), our study shows that DNA damage-primed macrophages exhibit an anti-inflammatory phenotype. This is consistent with a study that showed that IL6 is SWI/SNF-dependent in LPS-stimulated macrophages, while it is SWI/SNF-independent in LPS-stimulated primary mouse embryonic fibroblasts (MEFs), thereby implying that IL6 is induced in a cell-type-specific manner by a given stimulus (46).

The function and polarization state of macrophages is regulated through specific transcriptional programs in a context-dependent manner. These gene expression programs have been attributed to epigenetic regulation at the level of histone modification and chromatin remodeling (47–49). This raises the question whether chromatin plasticity can be viewed as multifaceted signal integration platform to regulate the phenotypic plasticity of macrophages. The ATP dependent chromatin remodelers, histone chaperons and histone modifiers play central role in transient disruption of chromatin organization post DNA damage (31–34).

Our phospho-proteomics data show differential regulation of phosphosites of ATP-dependent chromatin remodelers such as SWI/SNF Complex and NuA4 complex, histone chaperones such as CAF-1 and CENPA specific chaperones and many histone modifying enzymes in response to UV exposure of macrophages (Figure 4). The histone chaperons promote new histone deposition (50) with DNA damage specific histone modifications (51). This could reshape the epigenetic landscape leading to modulation of gene expression thus reprogramming the macrophage and altering its identity in response to genotoxic stress (31,52).

The functional assays showed that after exposure to UV or chemotherapeutics drugs, macrophages react differently to LPS and *E.coli* bioparticles. Interestingly, these functional adaptations were lost upon the treatment of M_DDR_ with the histone deacetylase inhibitor TSA or an ATM inhibitor. The data indicate that DNA damage-induced ATM mediates chromatin remodeling and epigenetic modification in macrophages, which acts as DNA damage memory and upon re-challenge, alters the functional response of macrophages. Therefore, these findings lend further credence to the notion that DNA damage scars persist as epigenetic memory and could potentially modulate macrophage function. The M_DDR_ macrophages not only adapt to exposure to genotoxic agents by acquiring phenotypic characteristics but also creates a DNA damage memory, modulating its response to subsequent immunogenic insult, a phenomenon analogous to immune memory.

Recently, the significant role of epigenetic regulation in macrophage biology and polarization has been more evident (53,54). Schmidt *et al*. have demonstrated that macrophages acquire specific epigenetic and transcriptional signature in response to various stimuli (55). Several reports have shown epigenetic regulation of macrophages in the context of infectious and chronic inflammatory diseases mostly involving M1s (49,53,56,57). In particular, histone methylation and acetylation adjacent to inflammation-related genes contribute to M1 or M2 phenotypes. Interestingly, histone methyltransferases are reported to be strongly associated with M2 activation by repressing M1 phenotype signature genes and promoting the transcription of M2 genes while histone demethyltransferase modulates polarization to both M1 and M2. Histone acetyltransferases are involved in initiating gene expression in macrophages during inflammation while histone deacetylases induce epigenetic changes to facilitate alternative gene expression (49). Accordingly, many pharmacologic modulators of epigenetic enzymes are documented to influence macrophage polarization (58). This raises the question of whether chromatin plasticity in macrophages can be viewed as a multifaceted signal integration platform for environmental cues.

The analysis of the differential PTMs indicated that the DDR including the regulation of the epigenetic changes is exerted by ATM. Indeed, we demonstrate that ATM functions as central regulator of DNA damage induced effects leading to the M_DDR_ phenotype. ATM is one of the apical kinases responsible for cellular responses to DNA damage including alterations in chromatin structure, DNA repair and transcription regulation (59–62). ATM is activated in response to DNA double strand breaks and also upon UV-induced DNA lesions (63). The ATM dependent DNA damage checkpoint temporarily arrest cell cycle or induces permanent cell cycle arrest that is senescence (64). Furthermore, ATM triggers chromatin alteration to facilitate accessibility of damaged chromatin to DNA repair proteins, followed by restoration of chromatin organization (31–34).

In accordance with our findings, Figueiredo et al. demonstrated that in response to anthracycline mediated DNA damage, ATM, ATR and Chek1 act as mediators of anti-inflammatory effects that can protect against sepsis (65). Additionally, recent studies reported elevated IL8 and IL6 levels (an inflammatory phenotype) in A-T patient serum (66,67). Our results are thus highly consistent with prominent role of ATM in regulating an anti-inflammatory response.

Interestingly, the immune checkpoint PD-L1 was also upregulated in macrophages primed with UV or DNA damage-inducing chemotherapeutic drugs. Our data indicate that chemotherapeutic drug treatment elicits “off-target” immunomodulatory effects such as an immunosuppressive phenotype and PD-L1 expression in macrophages. In agreement, recent studies have also shown that combining radiotherapy and chemotherapy with an immune checkpoint (PD-1 and PD-L1) inhibitor treatment significantly improved the survival of cancer patients (68,69). Moreover, targeting DDR proteins increased PD-L1 expression, thus enabling the anti-tumor response of antibodies against PD-L1 in SCLC and breast cancer models (70,71). The PD-L1 expression is positively correlated with γH2A.X (72) and it is upregulated in ATM/ATR/Chk1 kinases-dependent manner in cancer cells in response to DSBs (73).

Our results link immunological responses to chemotherapy and DDR with immune checkpoint therapy in macrophages. Targeting the DDR in macrophages during cancer therapy represents a novel strategy for enhanced macrophage mediated clearance of cancer cells. In this context, combining chemotherapy with immune checkpoint inhibitor as a front-line combinatorial principle appears as a novel rationale that was recently adopted in clinical routine with significantly improved survival of cancer patients such as in non-small lung cancer first line treatment with pembrolizumab plus chemotherapy (Deng et al., 2014; Twyman-Saint Victor et al., 2015);Gandhi et al.2018).

Finally, our findings allocate a central regulatory role for macrophages in the tumor microenvironment in the response to DNA damage-based therapies. The M_DDR_ phenotype represents an immune suppressive element capable of removing cellular debris and simultaneously limiting adaptive T-cell responses by PD-L1 induction. These findings offer a mechanistic rationale for the recent clinical results. In combining conventional chemotherapy with checkpoint inhibitors and targeted drugs will improve the understanding and development of combinatorial approaches in cancer therapy.

## Material and Methods

### Cell culture

The murine macrophage cell lines J774A.1, RAW264.7 were cultured in DMEM (Gibco) supplemented with 10 % FBS (Biochrom GmbH) and 1% Pen/Strep (Gibco). THP-1 cells were cultured in RPMI 1640 containing 10 % FBS and 1 % Pen /Strep. The THP-1 monocytes (10^6^ cells/ml) were plated in a 100 mm Petri dish (Sarstedt) and treated with 100μM PMA for 24 h. All cells were kept at 37°C in a 5% CO2 incubator (Binder). The cells were split every 2-4 days, at a confluence of 70-90 %. Peritoneal macrophages were cultured in DMEM (Gibco) supplemented with 10 % FBS (Biochrom GmbH) and 1% Pen/Strep (Gibco). The J774A.1 and RAW264.7 cells were scraped off the cell culture Petri dish while peritoneal macrophages and PMA-treated THP-1 cells were trypsinized (0.5 % trypsin/EDTA (Gibco)) at 37° C. The trypsinization was stopped after 2-3 minutes by adding respective complete media. The live cell number was determined using trypan blue (Gibco) and a hemocytometer (Marienfeld).

The murine macrophage cell line J774A.1 and HEK293T derived ecotropic Phoenix cells were cultured in Dulbecco□s Modified Eagle Medium (DMEM) containing 10% fetal bovine serum (FBS) and 1% Penicellin/Streptomycin (P/S). The human-MYC/BCL2 (hMB) cell line (strain 102), generated by Leskov et al., was cultured in B Cell Culture medium composed of a 1:1 ratio of Iscove□s Modified Dulbecco□s Medium (IMDM) and DMEM, supplemented with 10% FBS, 1% P/S, 1% GlutaMAX and 1% β-Mercaptoethanol (74).

For generation of RNAi targeted J774.A1 macrophages the MLS and MLP retroviral vector system was used as previously published. Specifically, shRNA target sequences for p53: 5"-CCACTACAAGTACATGTGTAA-3” and for p38: 5"-ATACCACGATCCTGATGATGAA-3” were used.

### Primary cell isolation and culture

CD3+ and CD14+ cells isolated from healthy donor PBMC using MACS positive selection magnetic beads (Miltenyi, Bergisch Gladbach, Germany) directly plated out at a concentration of 5×10^5^ cells/ml in white 96-well plates (100μl/well) in RPMI 1640 medium + 10% FBS + 1 % P/S. Subsequently, Mafosfamide or Doxorubicin was added in a dilution series ranging from 0.33nM to 333.33μM (for Mafosfamide) or from 0.33nM to 500μM (for Doxorubicin) in triplicate for each concentration (Figure 6). Furthermore, three wells contained only medium (no cells), three wells did not receive a chemotherapeutic treatment and three wells were treated with dimethyl sulfoxide (DMSO) as control. The outer most wells were only filled with sterile PBS to reduce the effect of evaporation. The plates were then incubated for 48h at 37°C with 5% CO2.

To isolate peritoneal macrophages, wild type C57BL/6 mice were injected intraperitoneal with thioglycolate and macrophages obtained via peritoneal lavage after 4 days. Bone-marrow-derived macrophages were obtained from C57BL/6 mice and differentiated *in vitro* using recombinant M-CSF.

CD3^+^ and CD14^+^ cells isolated from healthy donor PBMC using MACS positive selection magnetic beads (Miltenyi, Bergisch Gladbach, Germany). Another part of the isolated CD14^+^ cells were then differentiated into macrophages using recombinant M-CSF (Miltenyi, Bergisch Gladbach, Germany). After harvest, the macrophages were plated out in a concentration of 5×10^5^ cells/ml in RPMI 1640 medium containing 10% FBS, 1% P/S and 50U/ml human recombinant M-CSF on a white 96 well cell culture plate. The same concentration series of Mafosfamide and Doxorubicin as for T cells and monocytes were added to the wells (Figure 6) and incubated for 48h at 37°C with 5% CO2.

For generation of bone marrow derived macrophages (BMDM) cells from the bone marrow, wild-type (wt) C57BL/6 mice were sacrificed by cervical dislocation and both femurs were extracted by removing all muscle and connective tissue from the bones. The bones were then cut open at the very ends and flushed with cold PBS by using a 27G needle to remove the bone marrow. The flushed bone marrow cells were thoroughly resuspended and filtered with a 100μm cell strainer before they were centrifuged at 1500rpm for 8 minutes. Remaining erythrocytes were lysed using 5ml ACK lysis buffer for 5 minutes at room temperature. After adding PBS, cells were again centrifuged like previously described. Per femur, cells were plated out in 10ml differentiation medium for 24h. The non-adherent cells were then counted and plated out at a concentration of 6×10^5^/ml on a 10cm bacterial dish in differentiation medium. After 48h, 4ml of fresh differentiation medium was added. After an additional 3 days, medium was replaced by normal DMEM + 10% FBS + 1% P/S and incubated for 24h. Cells were then scraped off, counted and plated out for experiments at a concentration of 5×10^5^ cells/ml. After plating out, cells were allowed to reattach and recover for 24h before the start of an experiment.

To isolate peritoneal macrophages, wild type C57BL/6 mice were injected intraperitoneal with thioglycollate (BD). After 4 days peritoneal lavage was performed. Upon resuspending the cells in DMEM +10% FBS +1% P/S, the cells were counted and plated out at and were allowed to reattach and recover for 24h before the start of an experiment.

### UV irradiation

Cells were irradiated with 254 nm UV-C light using Phillips UV6 bulbs. The irradiance was measured using a UVX digital radiometer and a UVX-25 probe from UVP before the irradiation every time. Media from the cells was aspirated, and they were subsequently washed with DPBS (Lonza) before irradiation. The Petri dish with cells was opened and placed under the UVC machine for the calculated time. Immediately after exposure cells were supplemented with complete fresh media and incubated at 37°C, 5% CO2.

### Chemical inhibitor and Chemotherapeutic treatment

Media was aspirated, and cells were subsequently washed with DPBS (Lonza) before irradiation. The Petri dish with cells was opened and placed under the UVC machine for the calculated time. Immediately after exposure, either fresh media with DMSO or fresh media with one of the inhibitors was added, or cells were incubated at 37°C, 5% CO2. The chemical inhibitors were used in following concentration. Trichostatin A (10 nM) (Sigma), KU55933 (10 μM) (Selleckchem).

The cells were washed with DPBS and replenished with fresh media supplemented with one of the drugs or DMSO (0.05 %). Subsequently, the cells were incubated overnight at 37°C and 5% CO2 incubator. The cells for experiments performed in the presence of chemical inhibitor were treated with supplemented with one of the chemotherapeutics drugs along with chemical inhibitor and incubated overnight at 37°C and 5% CO2 incubator. The chemotherapeutic drugs were used in following concentration: Doxorubicin (5μM) (Tocris Bioscience), Mafosfamide (5μM) (Santa Cruz), Etoposide (5μM) (Sigma).

### Endotoxin tolerance assay

The cells were primed with UV or chemotherapeutics according to the protocol mentioned above. After priming treatment, cells were incubated overnight (16 h) at 37°C and 5% CO2. The next day, cells were counted, and cells were plated at conc. of 10^6^ cells/ml/well in a 12-well plate with fresh media or with fresh media supplemented with 1000 ng/ml LPS (Sigma) for 6 hrs. After 6 h of LPS exposure (referred as 0 h time point), immediately supernatants were collected. Levels of IL6 and TNFα were assessed with a murine and human ELISA kit (R&D Systems) respectively, according to the company’+s protocol. For qPCR, cells were pelleted and frozen at 0 h.

### Polybead Phagocytosis Assays

Polybead Amino Microsphere 3μm latex beads were labeled with the fluorescent dyes CypHer5E or DyLight680, both mono-N-hydroxysuccinimide (NHS) ester-activated. The lyophilized powder was dissolved in 0.1M sodium carbonate (pH: 9,0) to a final concentration of 100mg/ml. Five-hundred microlitres of the latex beads were centrifuged at 3000xg for 5 minutes and the pellet was resuspended in 500μl 0,1M sodium carbonate (pH: 9,0). Three-hundred micrograms of CypHer5E or DyLight680 NHS ester was added and incubated for 2h at room temperature while rotated. Afterwards, the labeled beads were centrifuged at 3000xg for 5 minutes and washed three times in 20ml PBS (with the same centrifugation conditions). Finally, beads were resuspended in 1ml PBS and correct labeling as well as the concentration of the beads was measured and calculated via flow cytometry. Labeled beads were stored at 4°C.

### pHrodo phagocytosis assays

Cells were primed according to the protocol mentioned above. The primed cells were counted, and 100,000 cells per well were plated in 96 well-plate. Cells were allowed to settle and adhere to the microplate for at least 1 h at 37°C 5% CO2. pHrodo® Red E.coli Bioparticles® (Life Technologies) were resuspended in 2 ml of Live Cell Imaging Solution (Cat. no. A14291DJ) and sonicated for 5 mins. The resuspended pHrodo® Red E.coli Bioparticles® were incubated with the cells for 1 h at 37 °C and then measured using PerkinElmer Enspire, fluorescence plate reader according to the company’+s protocol.

### Bead based phagocytosis assay

J774A.1 cells (wt as well as KDs) were plated out as follows: 1×10^5^ cells per 12 well for DMSO and negative control and 2×105 cells per 12-well for treated samples. Murine BMDM and human M-CSF differentiated macrophages were plated out at a concentration of 5×10^5^ cells per 12 well for all conditions. After 6h of reattachment for J774A.1 cells and 24h for primary macrophages, cells were treated with 5μM Mafosfamide, Doxorubicin, Nutlin-3A or the same volume of DMSO (0.05%) for 24h. Then, the medium was changed to fresh medium, containing 5×10^5^ DyLight680 labeled latex beads. The plate was centrifuged at 300xg for 1 minute to promote interaction of beads and cells. The incubation time of beads and macrophages was further optimized from 16h to 1.5h due to a very rapid phagocytosis of the beads by macrophages in general. After this time, non-phagocytosed beads were washed off the plate two times with PBS and cells were scraped off the plate and transferred to a 5ml FACS tube. Cells were centrifuged at 300xg for 5 minutes at 4°C, resuspended in 100μl PBS and stained with 1μl of CD11b-BrilliantViolet421 antibody for 20 minutes at 4°C. After 2 washes with PBS and centrifugation at 300xg for 5 minutes, the cells were resuspended in 250μl PBS and 50μl of this cell suspension were measured with the MACSQuant X flow cytometer. Analysis was performed with the MACSQuant software. Cells were gated for living and single cells, before the amount of double positive cells in the V1 (BrilliantViolet421) and R2 (DyLight680) channel was determined. For the transduced J774A.1 KD cell lines, single cells were also gated for GFP+ cells and only these were further used for the identification of double positive cells. The percentage of phagocytic active macrophages was normalized to each DMSO control. The ratio between counted cells via flow cytometry and used beads was calculated for each sample.

### Cell-based phagocytosis assay

On a 12 well plate, 2×105 J774A.1 wt and KD cells were plated out in 1ml DMEM + 10% FBS + 1% P/S. After 12h of reattachment, they were treated with 5μM Mafosfamide, Doxorubicin or Nutlin-3A for 24h. Treated cells were then scraped off the plate, counted and 4×104 cells per well in 100μl DMEM + 10% FBS + 1% P/S were seeded on a 96 well plate. Per condition, 5 replicates were seeded. Additionally, 10 wells were only filled with 100μl medium for hMB single cell control. The outer most wells were only filled with PBS to reduce evaporation. After 6h of reattachment, 1.5×105 hMB cells in 100μl B cell medium were added to the macrophages and the wells that only contained medium. The coculture was incubated for 16h and the remaining GFP+ hMB cells in the medium were measured using the MACSQuant VYB flow cytometer. The percentage of phagocytosed hMB cells (difference of remaining cells to hMB single cell controls) was normalized to each DMSO control.

### Immunofluorescence staining

18mm glass coverslips (Menzel-gläser) were coated with Poly-L-lysine solution (Sigma-Aldrich, P4707) for 1 h at 37°C. After incubation, the coverslip was washed with DPBS and dried. Cells were plated onto coverslips and treated as per experiment. After 24 h of treatment, cells were fixed with 4% paraformaldehyde (Roth) for 10 mins. The cells were then washed twice with 1 x wash buffer (1 x DPBS containing 0.05% Tween-20) and permeabilized with 0.1% Triton X-100 for 5 mins. Next, cells were washed twice and blocked for 30 min in 1% BSA (Sigma). For SAHF foci staining, cells were mounted on a slide using a Fluoromount-G ™, with DAPI (Invitrogen ™, 00-4959-52). For morphology imaging, cells were incubated with TRITC/conjugated Phalloidin (1:1000), and FITC conjugated secondary antibody for 60 minutes. The cells were rewashed thrice and mounted onto a slide using Fluoromount-G ™, with DAPI. The images were visualized with a confocal fluorescent microscope.

### Quantitative RT-qPCR

The RNA was extracted using RNeasy mini kit (Qiagen). The RNA was quantified using a Nanodrop 8000 spectrophotometer (Thermo Scientific). 1 μg of extracted RNA was dissolved in RNase-free water to get a total volume of 11.5 μl, and then the reverse transcription (RT) reaction was carried out. The RNA sample was denatured by incubating it at 70°C for 2 min in S1000TM or C1000TM Thermal Cycler PCR machine (Bio-Rad). The denatured RNA was quickly transferred on ice, and 8.5 ml of RT reaction Master-mix containing Superscript III (Invitrogen) was added. The PCR mix was placed back in the PCR machine to complete the RT PCR run for generating the complementary DNA (cDNA). Then, the cDNA sample was diluted by adding 80 μl distilled water and stored at −20 C. The cDNA samples generated with RT PCR were then used for performing qPCR using SYBR Green I (Sigma) and Platinum Taq polymerase (Invitrogen) on CFX96TM Real-Time PCR Detection System (Bio-Rad). The cDNA was added to qPCR buffer Master-mix, and 20 μl of it was pipetted per well in the 96-well plate (BIOplastics), and the corresponding 5 μl of Primer Master-mix was added followed by sealing the plate with Microseal ® adhesive sealer (Bio-Rad). The 60S acidic ribosomal protein P0 (rplp0), Peptidyl-prolyl cis-trans isomerasa (ppia) and Beta-2 microglobulin (β2m) were used as internal control to which the expression levels were normalized. The sequences of primers used in this study are as follows:

Il6 fwd ACACATGTTCTCTGGGAAATC, Il6 rev AAGTGCATCATCGTTGTTCATACA;

cd206 fwd TGCCGACATGCCAGGACGAAA, cd206 rev GTGGGCTCTGGTGGGCGAGT;

rplp0 fwd TGAAATTCTGAGTGATGTGC, rplp0 rev TTGTACCCATTGATGATGGAG;

pipa fwd CAAGACTGAATGGCTGGATG, pipa rev GTGATCTTCTTGCTGGTCTT;

β2m fwd CCCTGGTCTTTCTGGTGCTT, β2m rev ATTTCAATGTGAGGCGGGTG

### Fluorescence-activated cell sorting (FACS)

The cells were mock treated or primed with UV or chemotherapeutic drugs (according to protocol mentioned above) and incubated for 24 hrs. Next, cells were washed and scraped with DPBS. The cells were pelleted by centrifugation at 300xg for 5 minutes. The pellet was resuspended in 100μl of 1:10 dilution of FcR blocking reagent (Miltenyi Biotec) in DPBS and incubated at 4°C for 10 minutes in sterile FACS tubes. Afterward, cells were incubated for 15 minutes at 4°C either with 1μl of rat IgG2b isotype antibody or PE/Cy7 anti-mouse CD274 (B7-H1, PD-L1) antibody (Bio Legend) except for unstained control cells. Cells were washed twice and resuspended in 200 μl DPBS. Mean Fluorescence Intensity (MFI) was measured using the MACSQuant X flow cytometer for at least 10,000 cells. The MFI was analyzed using the MACSQuantify software. MFI values were isotype corrected and normalized to the mock-treated cells.

### Phosphoproteomics sample preparation and data acquisition and processing

The J774A.1 cells were treated with 10 J/m2 of UVC dose and incubated at 37° C, 5%CO2. The samples were collected after 4 h of UV exposure. CECAD / ZMMK Proteomics Facility carried out the sample preparation, data acquisition, and processing. In brief, 1.5 mg protein/sample was lysed in 8M Urea And digested in-solution using Trypsin and Lys-C. Next, generated peptides were desalted using 50 mg C18 Sep Pack-columns (Waters, #WAT 054955). Phosphopeptides were enriched using the High-Select™ TiO2 Phosphopeptide Enrichment Kit (A32993, Thermo Fisher) following manufacturer’+s instructions. All samples were analyzed on a Q Exactive Plus Orbitrap (Thermo Scientific) mass spectrometer that was coupled to an EASY nLC LC (Thermo Scientific). Phosphopeptide raw data were processed with Maxquant (version 1.5.3.8) using default parameters. Whole Proteome DIA data were processed and quantified using Spectronaut 10 (Biognosys).

The data and details about the used mass spectrometry settings are available via ProteomeXchange with identifier PXD013938. Reviewer account details:

#### Password

UcBG5wF4The softwares used for the phosphoproteome data analysis and representation are Cytoscape, Reactome database and InstantClue.

### Senescence β-Galactosidase staining

The cells were mock treated or primed with UV with and without chemical inhibitor (according to protocol mentioned above). After incubation at 37 °C and 5% CO2 for 24 h, samples were processed according to the company’+s protocol of Senescence β-Galactosidase staining kit (Cell Signaling Technology, 9860). The growth media was removed, and cells were rinsed with 2 ml of DPBS. Then cells were fixed by adding 1 ml of 1X Fixative Solution for 12 mins at room temperature. Next, the cells were washed 2 times with DPBS, and 1 ml of the β-Galactosidase staining solution was added to each well. The plate was then incubated overnight at 37°C, and cells were imaged using the EVOS FL Auto 2 imaging system.

### Annexin V staining

The cells were mock treated or primed with UV with and without chemical inhibitor (according to protocol mentioned above). After incubation at 37°C and 5% CO2 for 24 h, cells were washed twice with cold DPBS and resuspended in 1X Annexin binding buffer (BD Pharmingen™, 556454) at a concentration of 1 x 106 cells/ml. The 100 μl of the resuspended solution was incubated with 5 µl of FITC Annexin V (BD Pharmingen™, 560931) in FACS tube for 15 min at RT (25°C) in the dark. Next, 400 µl of 1X Binding Buffer was added to each tube, and cells were analyzed by flow cytometry within 1 hr.

### Statistics

Data were evaluated, and graphs were generated using GraphPad Prism 5. Unless otherwise stated, values represent the mean ± SD of three biological replicates. Statistical comparison of two groups was performed using an unpaired Students’+ two-tailed t-test. Differences were considered statistically significant at p-values less than 0.05 (ns; P > 0.05, P > 0.05; *, P ≤ 0.05; **, P ≤ 0.01; ***, P ≤ 0.001). All statistical analyses were performed using GraphPad Prism version 5.00 (GraphPad Software, San Diego, CA).

## Acknowledgments

We would like to thank Reinhild Brinker, Michael Michalik for excellent technical support and the CECAD proteomics facility. CP was supported by the “NRW -Nachwuchsgruppenprogramm” and Deutsche Forschungsgemeinschaft KFO286. BS acknowledges funding from the Deutsche Forschungsgemeinschaft (SCHU 2494/3-1, SCHU 2494/7-1, CECAD, SFB 829, SFB 670, KFO 286 and KFO 329), and the Deutsche Krebshilfe (70112899).

The authors declare no competing interests.

## Supplemental Figure Legends

**Supplementary Figure 1.** A micrograph example of breast cancer tissue (Giemsa) before (A-C) and after (D-F) neoadjuvant treatment displaying multiple macrophage with induction of PD-L1 post CTX treatment.

**Supplementary Figure 2. DNA damage modulates macrophage inflammatory response to LPS and phagocytosis** ELISA for IL6 quantification, supernatant collected from M_DDR_ and M_C_ macrophages at 0 h after LPS exposure in RAW264.7 (A), and peritoneal macrophages (C). ELISA for TNFα quantification, supernatant collected from M_DDR_ and M_C_ macrophages at 0 h after LPS exposure PMA-treated THP-1 (B). D) ELISA for IL6 quantification, supernatant collected from doxorubicin, mafosfamide and etoposide primed M_DDR_ and M_C_ macrophages 0 h-post LPS exposure. E) Relative fluorescence unit (RFU) is measured 1 h after M_DDR_ and M_C_ incubation with E.coli bioparticles called pHrodo in RAW264.7, THP-1, and peritoneal macrophages. Values represent the mean ± SD of three biological replicates \**P* < 0.05, \*\**P* ≤ 0.01 and \*\*\**P* < 0.001 (unpaired two-sided Student’+s *t*-test).

**Supplementary Figure 3.** A) The Log_2_ Fold change for the detected proteome B)PCA analysis variance ratio (C) for phosphoproteome. D) Correlation matrix with the Pearson correlation coefficient for phosphoproteome between 5 control (M_C_) and 5 UV primed (M_DDR_) replicates. Clustering was performed using Euclidean distance, complete linkage method. E) The distribution of phosphoproteome data. The phosphosites detected were 9160, out of which 1305 were significantly regulated, The significantly upregulated phosphosites were 854 and downregulated phosphosites were 451.

**Supplementary Figure 4. M**_**DDR**_ **phenotpe depends on ATM and p38 signaling** A) Induction of apoptosis assessed by Annexin V staining in J774.A1 macrophages treated ATM inhibitor KU55366, Doxorubicin, mafosfamide, etoposide and respective combinations B) Bead phagocytosis in J7774.A1 macrophages transduced with empty vector (MLS/MLP) or shRNA targeting p38 or p53 respectively. C) ADCP assessed in p53- and p38-shRNA targeted J774.A1 macrophages, RNAi mediated knock-down of p38 disrupts mafosfamide-induced increase of ADCP Values represent the mean ± SD of three biological replicates \**P* < 0.05, \*\**P* ≤ 0.01 and \*\*\**P* ≤ 0.001 (unpaired two-sided Student’+s *t*-test).

## Supplementary table legend

Table S1: The Whole Proteome DIA data processed and quantified using Spectronaut 10 (Biognosys) in five replicates of Control and UV treated J774A.1 macrophage cell line.

Table S2: The phosphoproteome data processed with Maxquant (version 1.5.3.8) using default parameters in five replicates of Control and UV treated J774A.1 macrophage cell line.

Table S3: The 1D enrichment analysis of annotated Gene Ontology (GO), Motifs, and Reactome terms. The yellow highlighted columns are represented in the figure 3B

Table S4: The 1D enrichment analysis showing overlap between enriched GO terms. The yellow highlighted columns are represented in the figure 3B

