## Supplementary figures and images for "ATM-mediated DNA damage response in macrophages primes phagocytosis and immune checkpoint regulation"

### Supplemental FIgures

##### Supplemental Figure 1

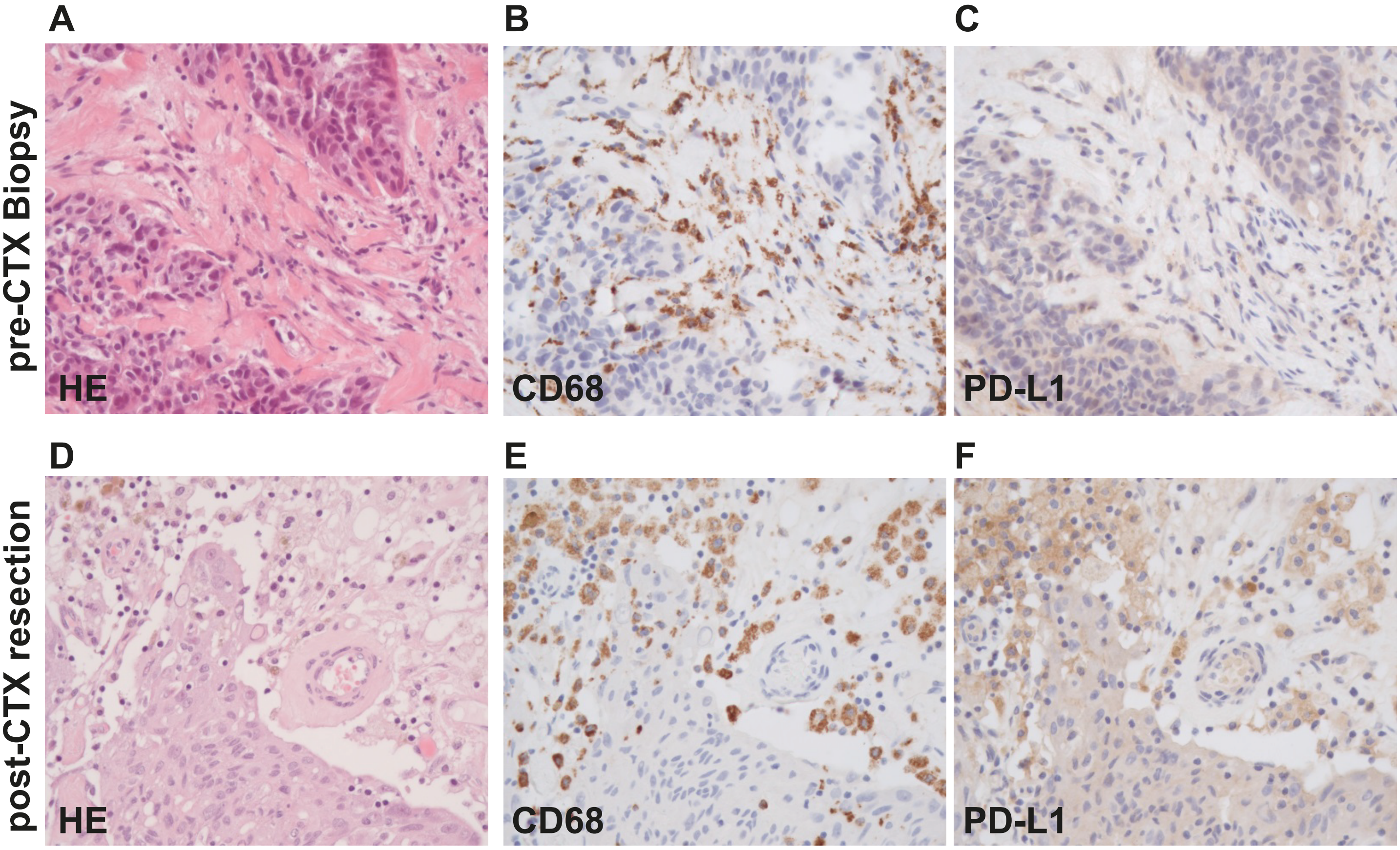

##### Supplemental Figure 2

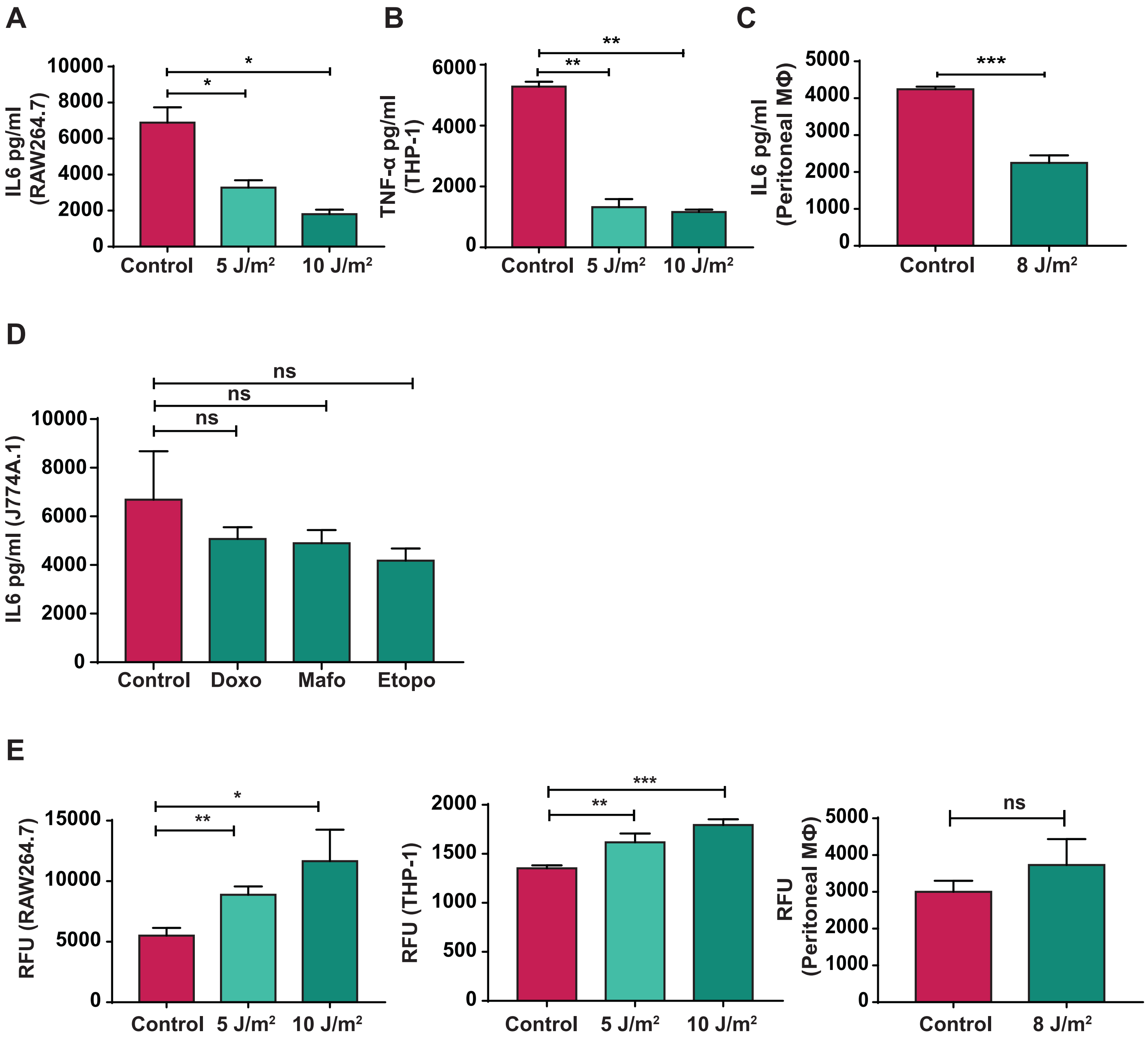

#### Supplemental Figure 3

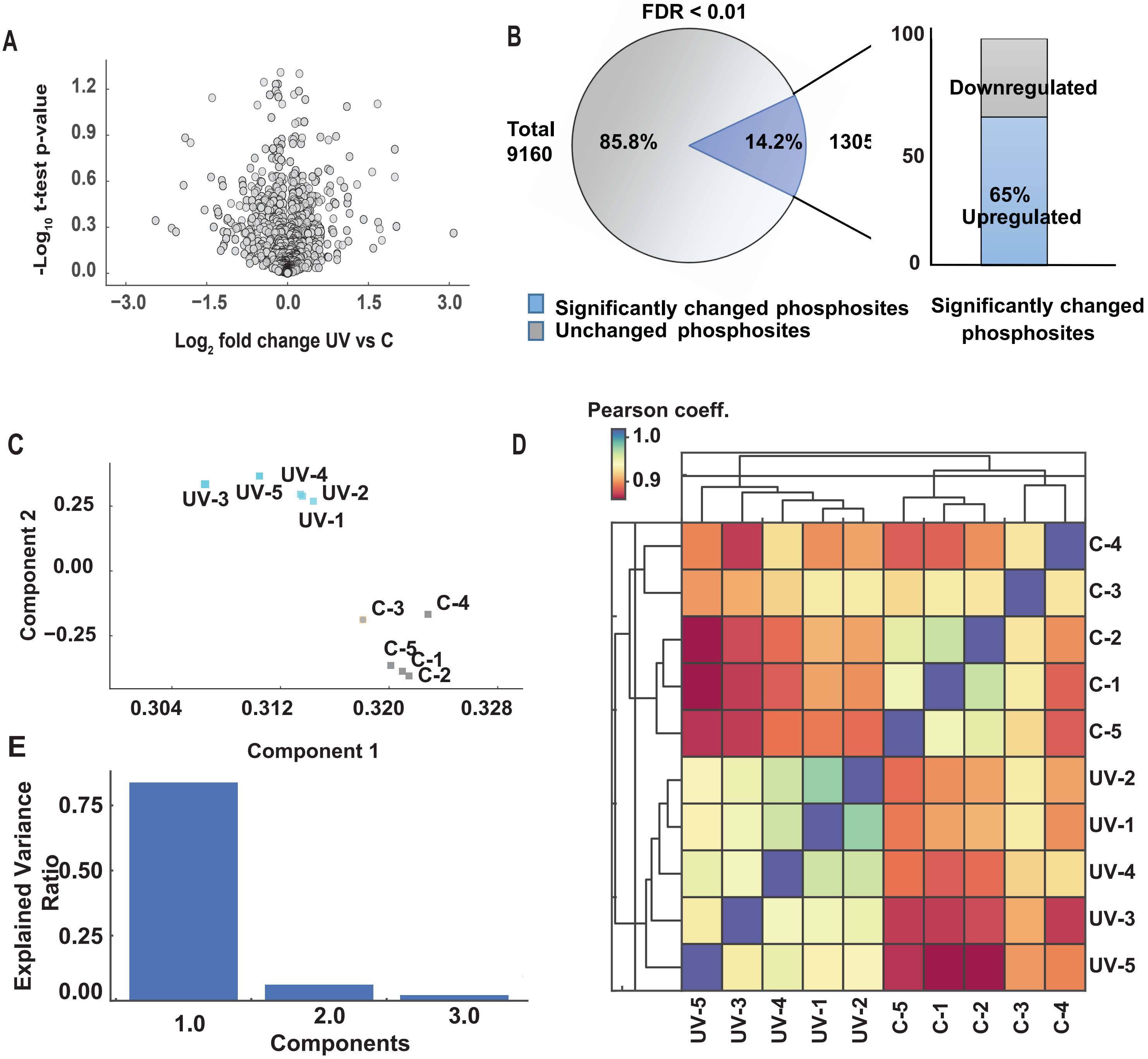

#### Supplemental Figure 4

## A

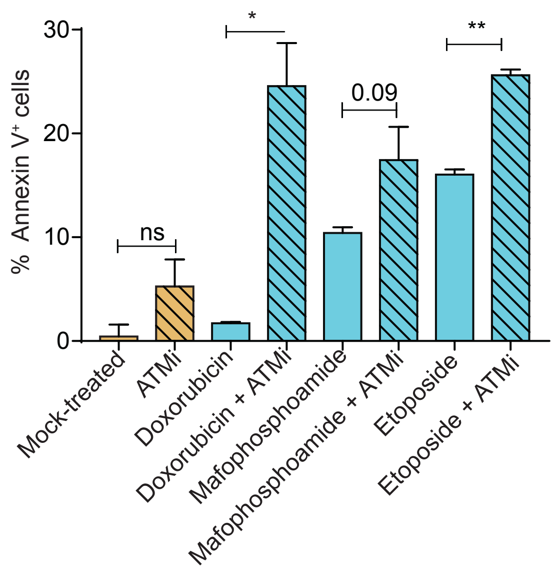


## B

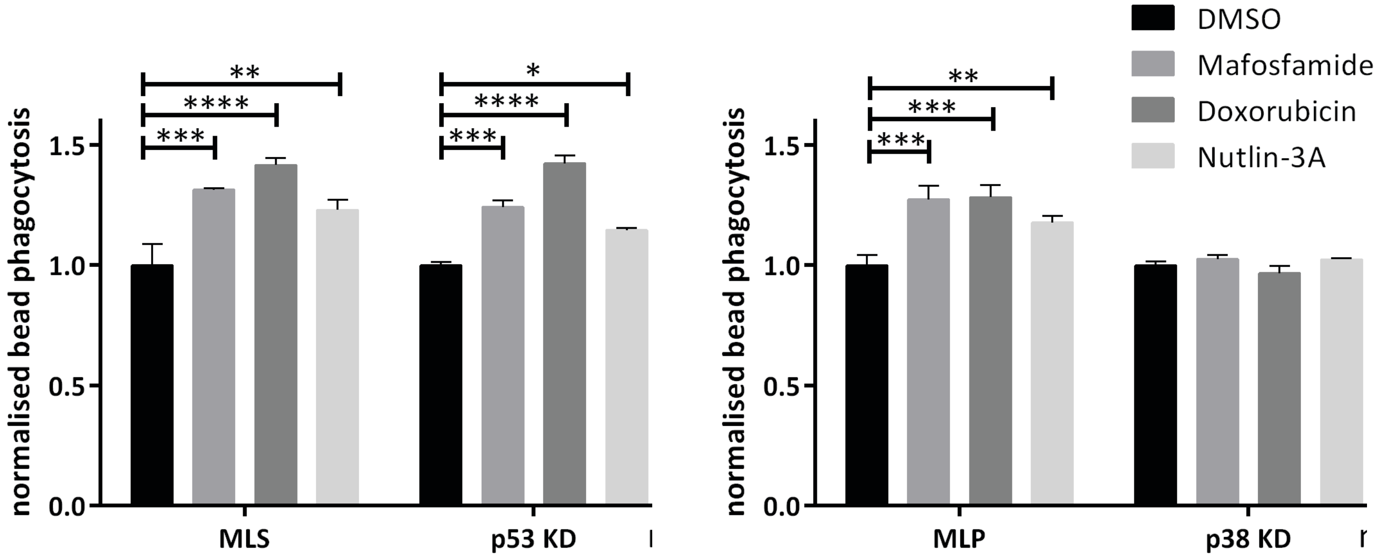


## C

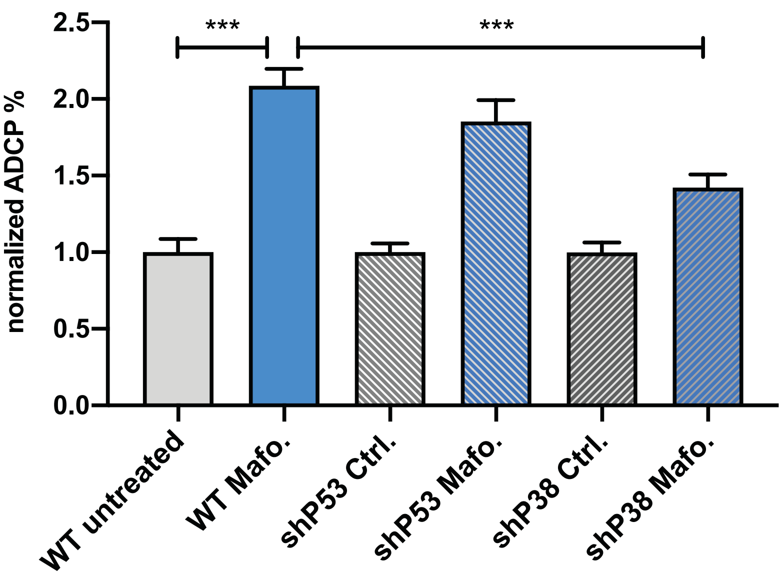
